# A ZBP1 isoform blocks ZBP1-mediated cell death

**DOI:** 10.1101/2024.02.02.578522

**Authors:** Zhi-Yu Cai, Puqi Wu, Wei Mo, Zhang-Hua Yang

**Author notes:** These authors contributed equally (Zhi-Yu Cai, Puqi Wu). Correspondence (W.M.); (Z.-H.Y.).

## Abstract

ZBP1 is an interferon-induced nucleic acid (NA) sensor that senses unusual Z-form NA (Z-NA), a type of left-handed nucleic acid. More than that, the binding of ZBP1 with Z-NA promotes cell death and inflammation. However, the mechanisms that dampen ZBP1 activation to fine-tune inflammatory responses are unclear. Here we characterize a short isoform of ZBP1 (referred to as ZBP1-S) as an intrinsic suppresser of the inflammatory signaling mediated by full-length ZBP1. Compared with ZBP1, ZBP1-S protein has Zα domains but no RHIM domains. Mechanistically, ZBP1-S depresses ZBP1-mediated cell death by competitive binding with Z-NA for Zα domains of ZBP1. Cells from mice (*Rip1^D^*^325^*^A/D^*^325^*^A^*) with Cleavage-resistant RIP1-induced autoinflammatory (CRIA) syndrome are alive but sensitive to IFN-induced and ZBP1-depedent cell death. Intriguingly, *Rip1^D^*^325^*^A/D^*^325^*^A^* cells go death spontaneously when ZBP1-S was deleted, indicating the cell death driven by ZPB1 is under the check of ZBP1-S. Thus, our findings reveal that alternative splicing of *Zbp1* represents an autogenic inhibition for regulating ZBP1 signaling and indicate that uncoupling of Z-NA with ZBP1 could be an effective strategy against auto-inflammations.

**Highlight:**

- ZBP1-short isoform is expressed synchronously with ZBP1.
- ZBP1-short isoform counteracts ZBP1 mediated cell death.
- ZBP1-S suppresses ZBP1 signaling in an Zα-domain dependent manner.
- ZBP1-S prevents the autoactivation of ZBP1 in *Rip1^D^*^325^*^A/D^*^325^*^A^* cells.

## Introduction

Nucleic acids (NAs), particularly double-stranded RNA (dsRNA) and dsDNA from pathogens, are detected by pattern recognition receptors to generate interferons (IFNs), which provide host defense against invading microbes^1^. Additionally, the sensors can also recognize endogenous NAs, leading to sustained and profound IFN production in various autoimmune diseases^2^. Typically, dsDNA and dsRNA favor to exist in a loose righthanded B form and a compact righthanded A form, respectively, under physiological conditions. In contrast to A and B, the Z conformation nucleic acids (Z-NAs) are a left-handed double helix with a zig-zag-shaped phosphodiester backbone that are energetically unstable^3^. Compared to right-handed DNA or RNA, left-handed Z-NAs emerges under cellular crisis such as heavy-duty viral infections and epigenome dysfunction, leading to grave inflammation and cell death^4,5^.

ZBP1 is the sensor of Z-NAs. Zα domains are expected to bind NAs in the Z conformation^6^. ZBP1 contains two N-terminal Zα domains, Zα1 and Zα2, followed by two receptor interacting protein homotypic interaction motifs (RHIMs) that have capacity to recruit RHIM proteins like RIP3 or RIP1^7^. Compared to other NA sensors, ZBP1 drives more robust inflammation via promoting cell death and should be well controlled. Upon Z-NA recognition, ZBP1 recruits RIP3 to initiate necroptosis, or promotes RIP1-FADD-caspase-8 complex to drive apoptosis^8^. As such, Z-NA-ZBP1 signaling axis plays vital roles in various pathological processes including influenza infection, embryonic lethality, and inflammatory bowel disease^4,9–12^.

By the Z-NA binding domain, viral protein E3 inhibits ZBP1-dependent cell death to escape viral proliferation restriction^13^. In mammal, another gene coding Zα domains is ADAR1 (Adenosine deaminases acting on RNA) which is an RNA editing deaminase rather than a Z-NA sensor^14^. Mutations in the sequence encoding the Zα domain of ADAR1 is associated with severe autoinflammatory disease, such as Aicardi-Goutieres syndrome (AGS)^15–19^. ADAR1 has been reported to suppress ZBP1-mediated cell death in several cell types and mouse models^20^. The depletion or Zα domain mutation of ADAR1 resulted in Z-RNA accumulation and activation of ZBP1, which culminated in RIP3-mediated cell death. Indeed, cell death in *Adar1^−/–^* cells and overt immunopathology caused by ADAR1 mutations in mice could be fully rescued by concurrent deletion of ZBP1 or the Zα2 domain of ZBP1. It has been well documented that ADAR1 regulates the metabolism of Z-NAs rather than directly blocks ZBP1 activation, which promotes us to find the endogenous Zα proteins that fine tune ZBP1-dependent inflammatory responses.

Here we reported an autogenic inhibition of ZBP1 activation. We identified a short isoform of ZBP1 (referred to as ZBP1-S), that only contains the two Zα domains, acts as a Z-NA competitor via its Zα domains to counteract ZBP1-dependent signaling in various cellular systems. Additionally, we revealed that cells harboring cleavage-resistant mutant RIP1 is hypersensitive to IFN-induced and ZBP1-dependent cell death, which is also under the check of ZBP1-S. Thus, identification of ZBP1-S as an intrinsic inhibitor of ZBP1 signaling should expand the understanding of the mechanism of fine-tuning ZBP1-related inflammatory responses.

## Results

### ZBP1-S is a more sensitive interferon stimulated gene than ZBP1-L

Two different cDNA variants have been identified for murine *Zbp1*, a long isoform and a short isoform. The long isoform of ZBP1 (hereafter, ZBP1-L) contains two N-terminal Zα domains and C-terminal RHIM domains, while the short isoform of ZBP1 (hereafter, ZBP1-S) only contains the two Zα domains (Figure 1A). In contrast to the well-known ZBP1-L, the characteristics of ZBP1-S remains unclear. We used western blotting to analyze the expression pattern of the two ZBP1 isoforms in multiple tissues from mice. As expected, ZBP1 was predominantly expressed in two protein isoforms in multiple tissues tested (ZBP1-L, ∼60 kDa and ZBP1-S, ∼25 kDa) (Figure 1B). Given *Zbp1* is an ISG (IFN-stimulated gene), next we analyzed the dynamic expression pattern of the transcripts of ZBP1-S and ZBP1-L in mouse embryonic fibroblasts (MEFs) (Figure 1C) and small intestinal epithelial organoids (Figure 1D) upon IFN-stimulation. ZBP1-S showed obviously higher and faster induction of transcript than ZBP1-L following IFN-stimulation (Figures. 1C and D), indicating that ZBP1-S is a more sensitive ISG isoform than ZBP1-L. Interestingly, we found that the protein level of them get out of step with their mRNA levels (Figure 1E). We speculated that the fast turn-over of ZBP1-S protein might result from protein degradation. Indeed, tentative exploration data showed that ZBP1-S has more shorter half-life periods than ZBP1-L (Figure 1F). And further analysis revealed that ZBP1-S undergone degradation by proteasome but not autophagy, as the proteasome inhibitor MG-132 but not the autophagy inhibitor Chloroquine (CQ) could efficiently increase the protein level of ZBP1-S either induced by IFN (Figures 1G and S1A) or ectopically expressed (Figures S1B and C). Considering these features of ZBP1-S, it implied that ZBP1-S might possess the potential to be a negative regulator of ZBP1 signaling.

**Figure 1.**
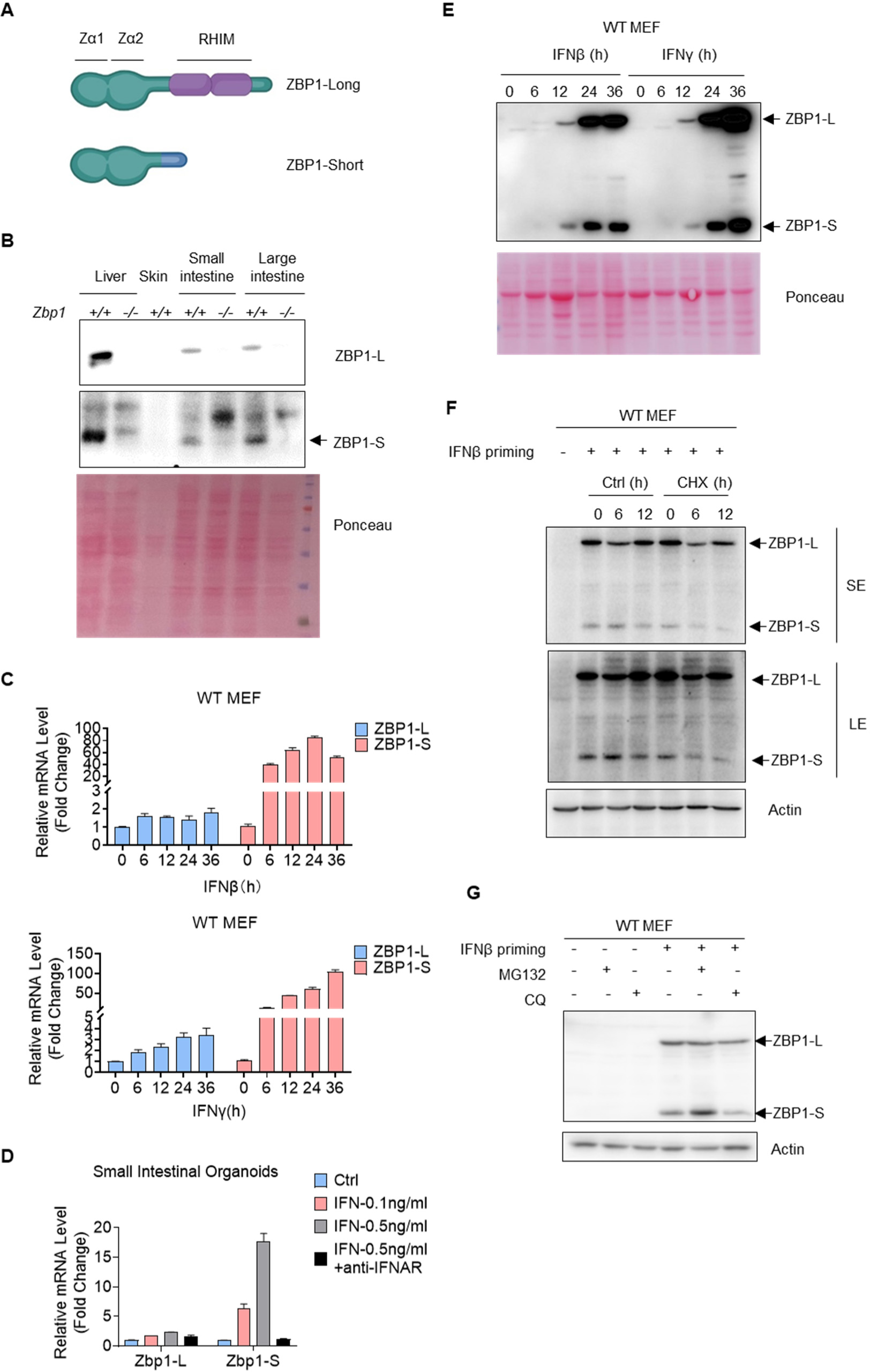
ZBP1-S is a more sensitive ISG than ZBP1-L. **A.** Graphic model for functional domains of ZBP1-S and ZBP1-L. **B.** Protein level of ZBP1-L and ZBP1-S detected by immunoblotting in liver, skin, small intestine, and large intestine. **C.** mRNA level change of Zbp1-L and Zbp1-S after stimulation of IFN β (10 ng/ml) and IFN γ (10 ng/ml) in MEFs. Timing of detection as indicated. Data are shown as mean ± s.e.m. of triplicates. **D.** mRNA level change of Zbp1-L and Zbp1-S in small intestinal organoids after stimulation of low concentration IFNβ (0.1 ng/ml), high concentration IFNβ (0.5 ng/ml) and high concentration IFNβ (0.5 ng/ml) with Anti-IFNAR antibodies for 12 h. Data are shown as mean ± s.e.m. of triplicates. **E.** Protein level of ZBP1-L and ZBP1-S detected by immunoblotting after stimulation of IFNβ (10 ng/ml) and IFNγ (10 ng/ml) in wild-type (WT) MEF cells. **F.** Protein half-life of ZBP1-L and ZBP1-S detected by immunoblotting after IFNβ (10 ng/ml) priming for 24 h and treated with Cycloheximide (CHX, 10 μg/ml) as indicated. **G.** WT MEF cells were treated with MG132 (5 μM) and CQ (10 μM) for 12 h after priming with IFNβ (10 ng/ml). Protein levels of ZBP1-L and ZBP1-S analyzed by immunoblotting.

### ZBP1-S inhibits ZBP1-dependent cell death

It has been well-documented that IAV-infection triggers ZBP1-dependent cell death^8^. To assess the role of ZBP1-S in ZBP1 signaling, we first overexpressed ZBP1-S in wildtype (WT) MEFs, which is sensitive to IAV infection. As indicated by PI staining (Figure 2A), overexpression of ZBP1-S efficiently inhibits necrotic cell death induced by PR8 and WSN, two commonly used IAV strains. Western blotting analysis revealed that both necroptosis (p-MLKL) and apoptosis (cl-PARP) induced by IAV were significantly attenuated by overexpressing ZBP1-S (Figure 2B). These data indicated that ZBP1-S might function as a negative regulator of ZBP1 signaling. To further confirm the negative role of ZBP1-S, we turned to assess the other contexts involving ZBP1 signaling. Previous study has shown that IFN-induced necroptosis in *Rip1* deficient cells was dependent on ZBP1^9,10,21^. Similar to IAV infection, overexpression of ZBP1-S significantly prevented necroptosis induced by IFNs in both *Rip1* knockout (KO) L929 cells and *Rip1^-/-^* MEF cells (Figures 2C-F and Figure S2A). RIP1 inhibits ZBP1-induced necroptosis by recruiting Casp8 and IFN-induced necroptosis in *Casp8* KO L929 is also dependent on ZBP1^21^. As expected, overexpression of ZBP1-S also significantly attenuated necroptosis induced by IFNβ in *Casp8* KO L929 cells (Figures S2B and S2C). Of note, overexpression of ZBP1-S had no effect on ZBP1-independent cell death, such as apoptosis-induced by TNF+Smac mimetic (TS) or staurosporine (STS), necroptosis induced by TNF+Smac mimetic+zVAD (TSZ), and ferroptosis induced by RSL3, indicating that ZBP1-S specifically inhibits ZBP1 signaling (Figure S2D). More Importantly, knock-down of ZBP1-S by siRNA induced spontaneous necroptosis under resting stage and also exacerbated necroptosis upon IFNs treatment in *Rip1^-/-^* MEF cells (Figures 2G-I). Collectively, these results demonstrated that ZBP1-S specifically suppressed ZBP1-mediated cell death.

**Figure 2.**
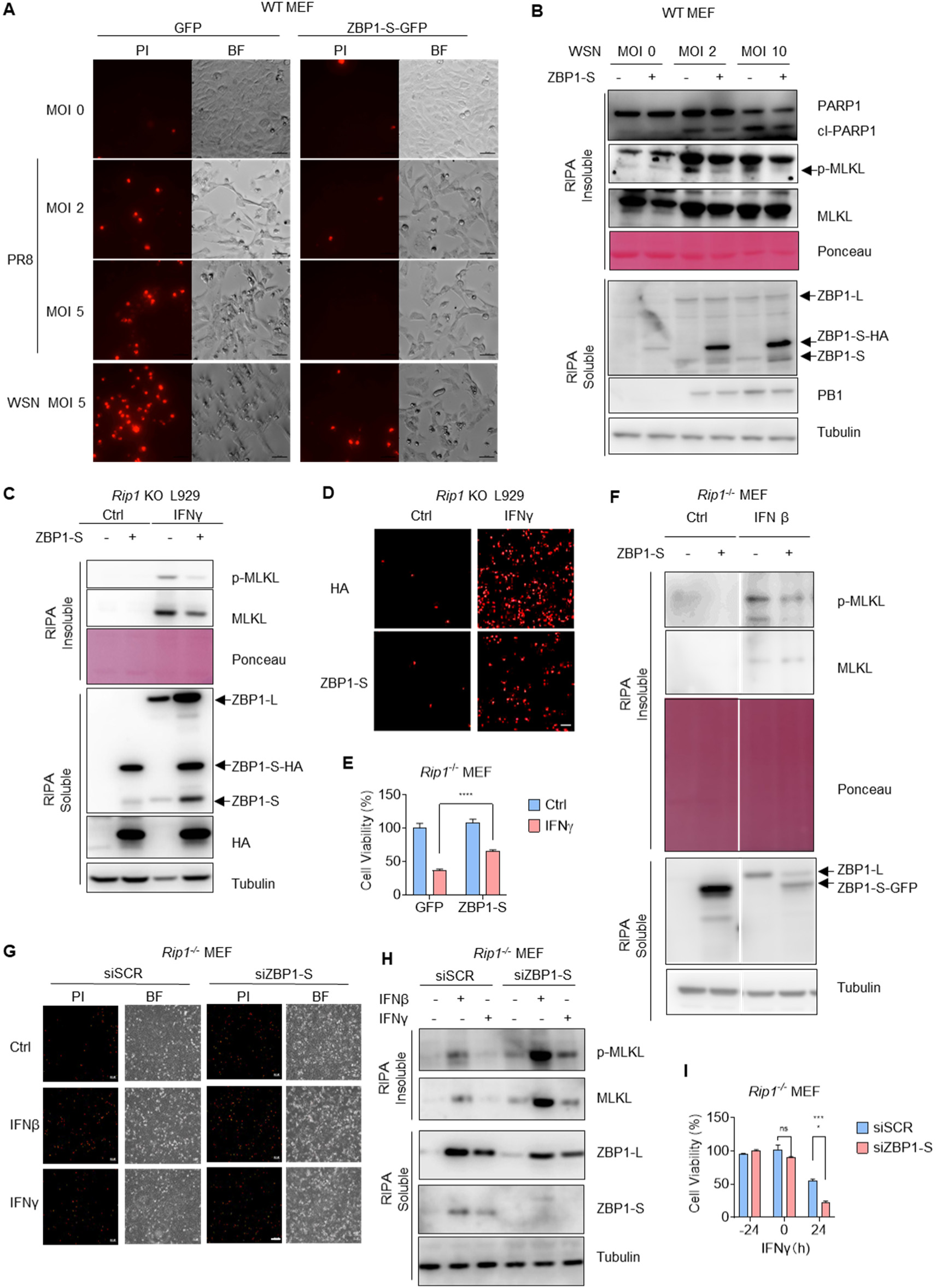
ZBP1-S suppresses ZBP1-dependent cell death. **A.** ZBP1-S-GFP and GFP were overexpressed in WT MEF cells. PI images were obtained after infection of influenza virus A for 30 h. The strain of Influenza A virus and the multiplicity of infection (MOI) were indicated. **B.** ZBP1-S-HA and HA were overexpressed in WT MEF cells. The cells were infected by WSN with the MOI as indicated for 24 h, and the cell lysates were analyzed by immunoblotting to detect the proteins as indicated. p-MLKL was detected in the RIPA insoluble fraction of cell lysates. Cl-PARP1, cleaved PARP1. p-MLKL, S345 phosphorylation of MLKL. **C.** ZBP1-S-HA and HA were overexpressed in *Rip1* KO L929 cells. The cells were treated by IFNγ (10 ng/ml) for 36 h. p-MLKL was detected in the RIPA insoluble fraction of cell lysates. **D.** The same cells in (C) were treated by IFNγ (10 ng/ml) for 36 h. PI images were obtained to detect cell death. Scale bar, 150 μm. **E.** ZBP1-S-GFP and GFP were overexpressed in *Rip1^-/-^*MEF cells. Cell survival (%) was measured after treatment of IFNγ (20 ng/ml) for 24 h. Data are shown as mean ± s.e.m. of triplicates.

**F.** The same cells in (E) were treated by IFNγ (20 ng/ml) for 24 h. The cell lysates were analyzed by immunoblotting to detect the proteins as indicated.

**G.** ZBP1-S was knocked down by siRNA in *Rip1^-/-^* MEF cells. PI images was obtained after treatment of IFNβ (20 ng/ml) and IFN γ (20 ng/ml) for 24 h.

**H.** The same cells in (G) were treated by IFNβ (20 ng/ml) and IFNγ (20 ng/ml) for 24 h. The cell lysates were analyzed by immunoblotting to detect the proteins as indicated.

**I.** Cell survival (%) of the same cells as in (G) was measured after treatment of IFNγ (20 ng/ml) for 24 h. Data are shown as mean ± s.e.m. of triplicates.

### ZBP-S suppressed ZBP1 signaling in a Z*α*-domain dependent manner

To further character the mechanism by which ZBP-S inhibits ZBP1-mediated cell death, we assessed the interaction between ZBP1 and ZBP1-S. We found that ZBP1 and ZBP1-S could interact with itself but could not interact with each other (Figures S3A-C). Compared to ZBP1, ZBP1-S contains two N-terminal Zα domain but loss the C-terminal RHIM domain, which links ZBP1 signaling to downstream RIP3, so we speculated that ZBP1-S might inhibits ZBP1 signaling by competing with ZBP1 for Z-NA binding. To test this possibility, we generated a Z-NA binding deficient ZBP1-S by mutating its two Zα domains (Zα1&Zα2 double mutant, ZαDM). We overexpressed WT and the ZαDM ZBP1-S into WT MEFs and found that in contrast to WT ZBP1-S, the mutant ZBP1-S could not inhibit IAV induced cell death anymore (Figures 3A-C and Figures S3D-E). Consistently, ZαDM ZBP1-S loss its ability to suppress IFN-induced necroptosis in either *Rip1* KO L929 cells (Figure 3D) or *Rip1* KO MEF cells (Figures 3E-G). Together, these results demonstrated that ZBP1-S suppressed ZBP1 signaling in a Zα-dependent manner.

**Figure 3.**
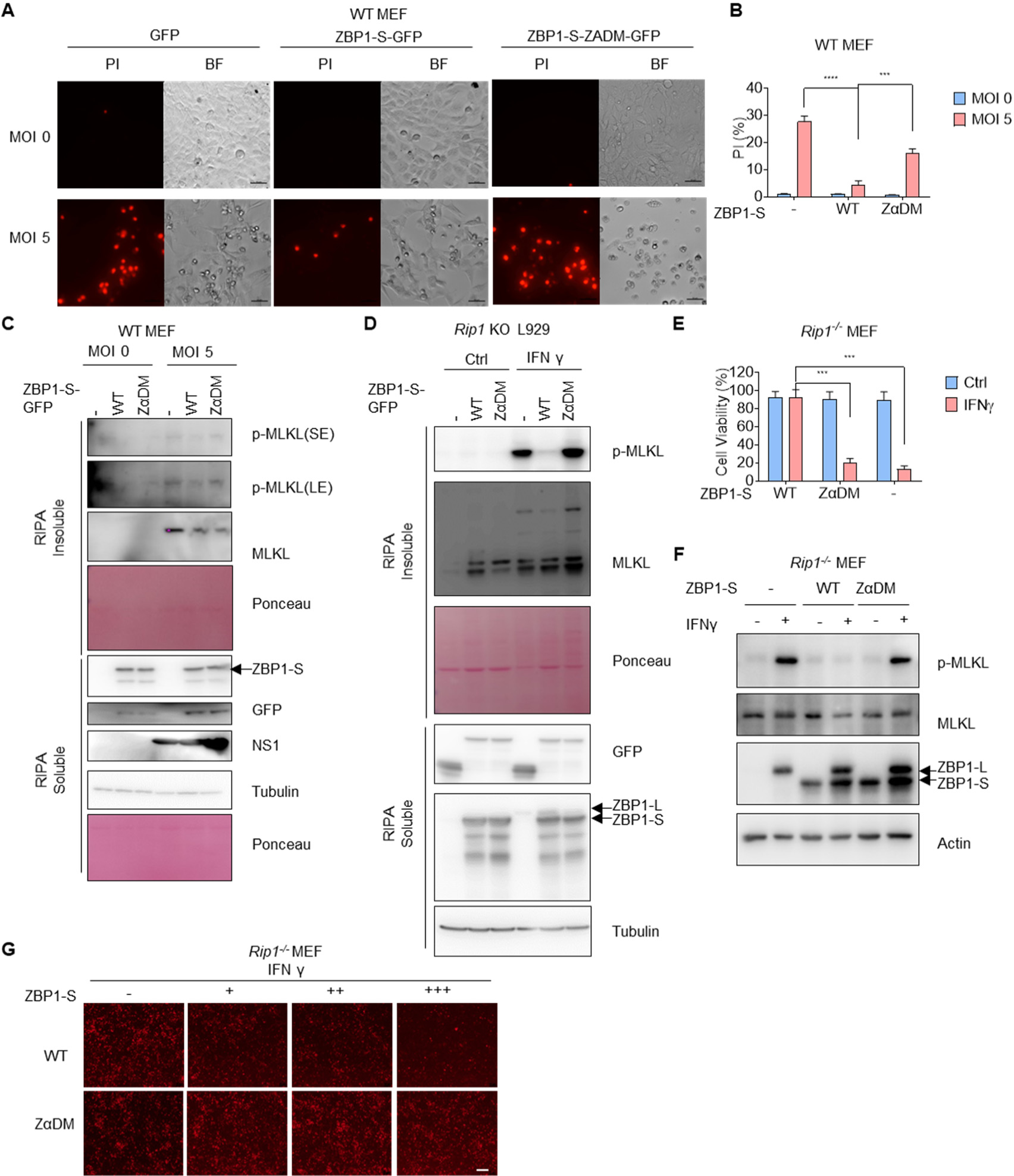
ZBP1-S inhibits ZBP1-dependent cell death by its Z*α* domain. **A.** GFP, ZBP1-S-GFP and ZBP1-S-ZαDM-GFP were overexpressed in WT MEF cells. PI images were obtained after infection of influenza virus A for 30 h. The strain of Influenza A virus and the MOI were indicated. ZBP1-S-ZαDM-GFP (Zα1& Zα1 domains Double Mutation), the mutation results in disability of Z-nucleic acid binding. Scale bar, 50 μm. **B.** Percentage of PI positive cells were measured from (A). Data are shown as mean ± s.e.m. of triplicates. **C.** The same cells in (A) were infected by influenza A virus for 30 h. The cell lysates were analyzed by immunoblotting to detect the proteins as indicated. **D.** GFP, ZBP1-S-GFP and ZBP1-S-ZαDM-GFP were overexpressed in *Rip1* KO L929 cells. The cells were treated by IFNγ (20 ng/ml) for 36 h. p-MLKL was detected in the RIPA insoluble fraction of cell lysates. **E.** GFP, ZBP1-S-GFP and ZBP1-S-ZαDM-GFP were overexpressed in *Rip1^-/-^* MEF cells. Cell survival (%) was measured after treatment of IFNγ (20 ng/ml) for 48 h. Data are shown as mean ± s.e.m. of triplicates. **F.** The same cells in (E) were treated by IFNγ (20 ng/ml) for 48 h. the cell lysates were analyzed by immunoblotting to detect the proteins as indicated. **G.** ZBP1-S-GFP and ZBP1-S-ZαDM-GFP were overexpressed in *Rip1^-/-^*MEF cells with increasing doses. PI images of the cells were obtained after treatment of IFNγ (20 ng/ml) for 48 h. Scale bar, 20 μm.

### ZBP1-S prevents the autoactivation of ZBP1 in *Rip1^D^*^325^*^A/D^*^325^*^A^* cells

Next, we measured the protective role of ZBP1-S under an autoinflammatory model. Mutations D324N, D324H and D324V in human RIP1 (corresponding residue is D325 in mice) have been found in rare familial human disease, named Cleavage-resistant RIP1-induced autoinflammatory (CRIA) syndrome^22–25^. These patients experienced recurring fevers along with lymphadenopathy. The patients showed strong RIP1-dependent activation of inflammatory signaling pathways with overproduction of inflammatory cytokines and chemokines. Blocking TNF cannot suppress inflammation in patients, suggesting that TNFα-RIP1 signaling is not the key pathogenic pathway to drive cell death and inflammation in CRIA patients^23,24^. IFNγ and Interleukin 6 (IL-6) were also detected markedly in the serum of CRIA patients. Then we generated MEFs from *Rip1^D^*^325^*^A/D^*^325^*^A^*mice which mimic the RIP1 cleavage-resistant mutation in CRIA patients to assess the pathogenic contribution of IFN-ZBP1 pathway. We examined the cell response by measuring cell survival with the CellTiter-Glo assay and assessed cell death by measuring the plasma membrane permeability with PI staining. We found that IL-6 treatment could not induce cell death in *Rip1^D^*^325^*^A/D^*^325^*^A^* MEFs (Figure S4A), while both IFNβ and IFNγ induced robust cell death in *Rip1^D^*^325^*^A/D^*^325^*^A^* MEFs but not wildtype (WT) MEFs (Figures 4A, B and Figure S4B). Western blotting analysis further revealed that the cell death induced by IFN in *Rip1^D^*^325^*^A/D^*^325^*^A^* MEFs was necroptosis and apoptosis as indicated by p-MLKL and cleaved Casp3 signals, respectively (Figure 4C). Besides cell death, IFNγ also dramatically increased the production of pro-inflammatory cytokines in *Rip1^D^*^325^*^A/D^*^325^*^A^* MEFs, like IL-6 (Figure S4C), TNFα (Figure S4D), and multiple innate-immune response related genes (Figure S4E). To confirm that the cell death induced by IFN was independent of the autocrine TNFα, the TNFα-neutralizing antibody was used. The data showed that TNFα-neutralizing antibody could not rescue the cell death induced by IFN in *Rip1^D^*^325^*^A/D^*^325^*^A^* cells (Figure S4F). In contrast, knock-down of ZBP1 nearly completely blocked IFN-induced cell death in *Rip1^D^*^325^*^A/D^*^325^*^A^*MEFs (Figure 4D), and significantly decreased the level of p-MLKL and p-RIP3 (Figure 4E). Thus, IFN-ZBP1 is indispensable for cell death and inflammation in *Rip1^D^*^325^*^A/D^*^325^*^A^*cells.

**Figure 4.**
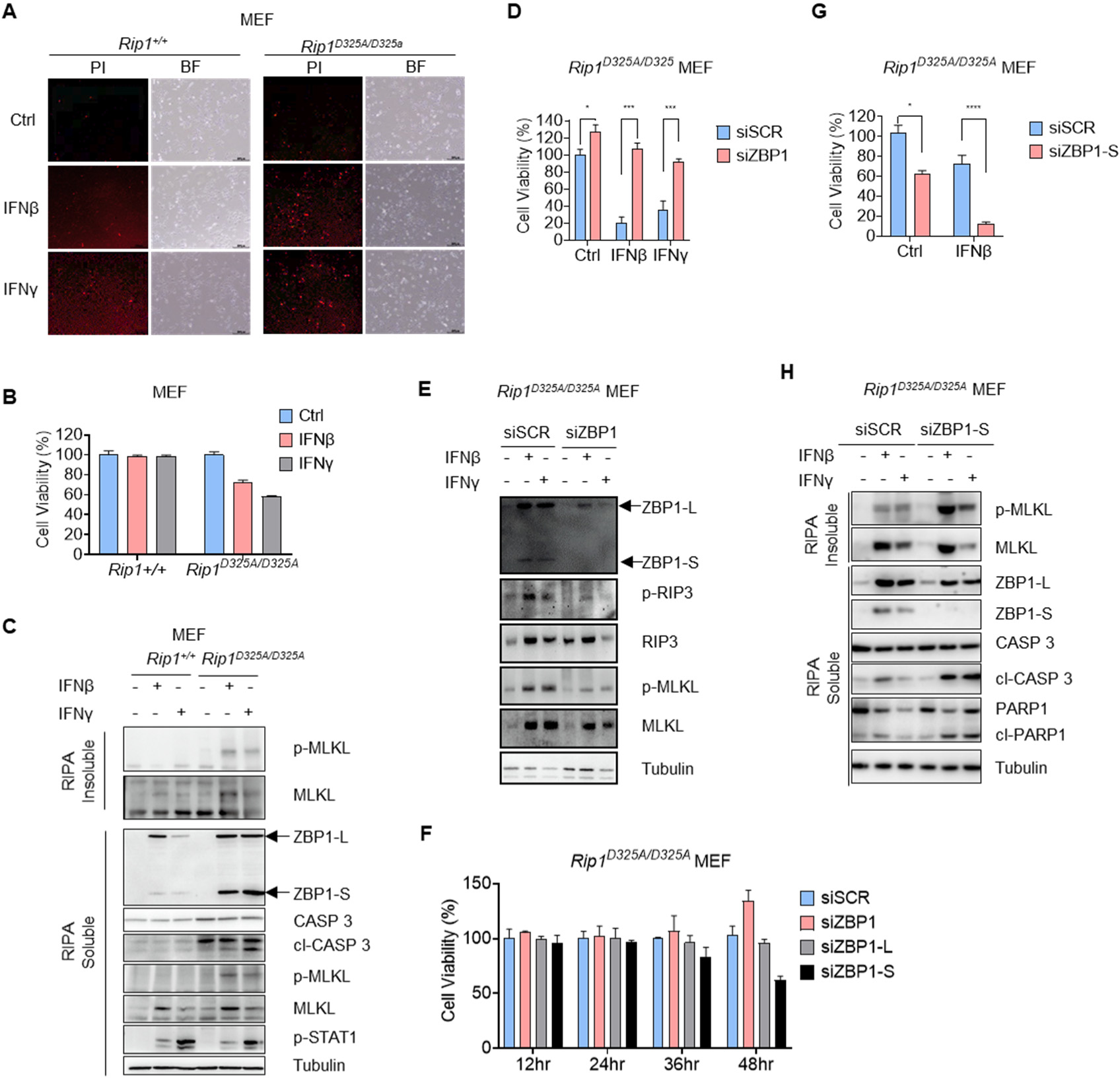
ZBP1-S prevents the autoactivation of ZBP1 in *Rip1^D^*^325^*^A/D^*^325^*^A^* cells. **A.** E1A+Ras-transformed *Rip1^D^*^325^*^A/D^*^325^*^A^* and *Rip1^+/+^*MEF cells were treated with IFNβ (20 ng/ml) and IFNγ (20 ng/ml) for 48 h. PI images were obtained to detect cell death. Scale bar, 150 μm. **B.** The same cells as in (A) were treated by IFNβ (20 ng/ml) and IFNγ (20 ng/ml) for 48 h. Cell survival (%) was measured. Data are shown as mean ± s.e.m. of triplicates. **C.** The same cells as in (A) were treated by IFNβ (20 ng/ml) and IFNγ (20 ng/ml) for 48 h. The cell lysates were analyzed by immunoblotting to detect the proteins as indicated. **D.** ZBP1 was knocked down by siRNA for 24 h in E1A+Ras-transformed *Rip1^D^*^325^*^A/D^*^325^*^A^* MEFs. Cell survival (%) was measured after treatment of IFNβ (20 ng/ml) and IFN γ (20 ng/ml) for 36 h. Data are shown as mean ± s.e.m. of triplicates. **E.** The same cells as in (D) were treated by IFNβ (20 ng/ml) and IFNγ (20 ng/ml) for 36 h. The cell lysates were analyzed by immunoblotting to detect the proteins as indicated. **F.** ZBP1, ZBP1-L and ZBP1-S were knocked down by siRNA for 24 h in E1A+Ras-transformed *Rip1^D^*^325^*^A/D^*^325^*^A^* and *Rip1^+/+^* MEFs. Cell survival (%) was measured at different period of time. Data are shown as mean ± s.e.m. of triplicates. **G.** ZBP1-S was knocked down by siRNA in E1A+Ras-transformed *Rip1^D^*^325^*^A/D^*^325^*^A^* MEFs. Cell survival (%) was measured after treatment of IFNβ (20 ng/ml) for 36 h. Data are shown as mean ± s.e.m. of triplicates. **H.** The same cells as in (G) were treated by IFNβ (20 ng/ml) for 36 h. The cell lysates were analyzed by immunoblotting to detect the proteins as indicated.

To further clarify the role of ZBP1 and ZBP1-S in IFN-induced cell death, we designed siRNA specifically targeting the two isoforms of ZBP1. We found that, knock-down of ZBP1-S resulted in the spontaneous cell death of *Rip1^D^*^325^*^A/D^*^325^*^A^* MEFs without stimulation of IFN (Figure 4F). Additionally, deficiency of ZBP1-S also dramatically exacerbated the cell death induced by IFN (Figures 4G, H and Figure S4G). Altogether, RIP1-D325A renders cells hypersensitive to IFN-induced ZBP1-dependent cell death and it is negatively regulated by ZBP1-S.

## Discussion

ZBP1 plays an essential role in innate immunity as a Z-NA sensor and a detonator of cell death. In this study, we identification an autogenic inhibition mechanism of ZBP1 activation. We reported that an isoform whose induction is more sensitive to interferon, ZBP1-S, inhibited ZBP1-mediated cell death by competing for Z-NA. Additionally, we revealed that IFN-induced cell death in cells containing cleavage-resistant mutant RIP1 is mediated by ZBP1, which is also under the check of ZBP1-S. During the study, we also found several phenomena worth further exploration. First, we found that ZBP1-S protein has a shorter half-life than ZBP1-L, and this faster turnover is mediated by proteasome. Further exploration is still needed to understand how the proteasome senses the induction of ZBP1-S to mediates the rapid degradation of ZBP1-S but not ZBP1 itself. Preliminary analysis showed that ZBP1-S is a highly intrinsically disordered protein, and we speculated that ZBP1-S may be rapidly degraded by ubiquitin-independent proteasome, like SE protein in plants and p53 in mammal^26,27^. The biological significance of rapid degradation of ZBP1-S protein by proteasome need further investigation. The rapid transcriptional induction of ZBP1-S enables it to respond promptly to the presence of Z-NA before the engagement of ZBP1-mediated cell death, while the rapid degradation of ZBP1-S protein by the proteasome may be necessary for the restoration of cell homeostasis. Second, although numerous studies have shown that cells harboring cleavage-resistant RIP1 are hypersensitive to TNF-induced cell death, we have now revealed that these cells are also sensitive to IFN-induced cell death. Whether this finding is true in vivo needs further investigation. Notably, despite the fact that most of our knowledge of RIP1 function comes from investigation of TNF signaling, and RIP1-D325A cells are hypersensitive to TNF stimulation, patients with CRIA syndrome responded poorly to TNF inhibitors. It will be interesting therefore to determine whether IFN neutralizing antibodies has effect in human CRIA syndrome. Finally, understanding the regulation of alternative splicing of ZBP1 could provide a new perspective for manipulation of ZBP1-related inflammatory diseases. Through targeting alternative splicing, it is possible to not only reduce the amount of ZBP1-L, but also increase the level of antagonistic ZBP1-S, potentially achieving a doubling effect on ZBP1-dependent cell death and inflammation.

## Acknowledgements

The study was supported by the National Natural Science Foundation of China (32225016 to W.M. and 32170751 to Z.-H.Y.), the National Key R&D Program of China (2021YFA1101401 to W.M.).

## Author contributions

Conceptualization and Supervision: Z.-Y.C., W.M. and Z.-H.Y.; Data curation: P.-Q.W., Z.-Y.C. and Z.-H.Y.; Formal analysis: Z.-Y.C., W.M. and Z.-H.Y.; Writing-original draft, Z.-Y.C., W.M. and Z.-H.Y.; Writing-review & editing: W.M. and Z.-H.Y.

## Declaration of interests

The authors declare no competing interests.

**Figure S1.**
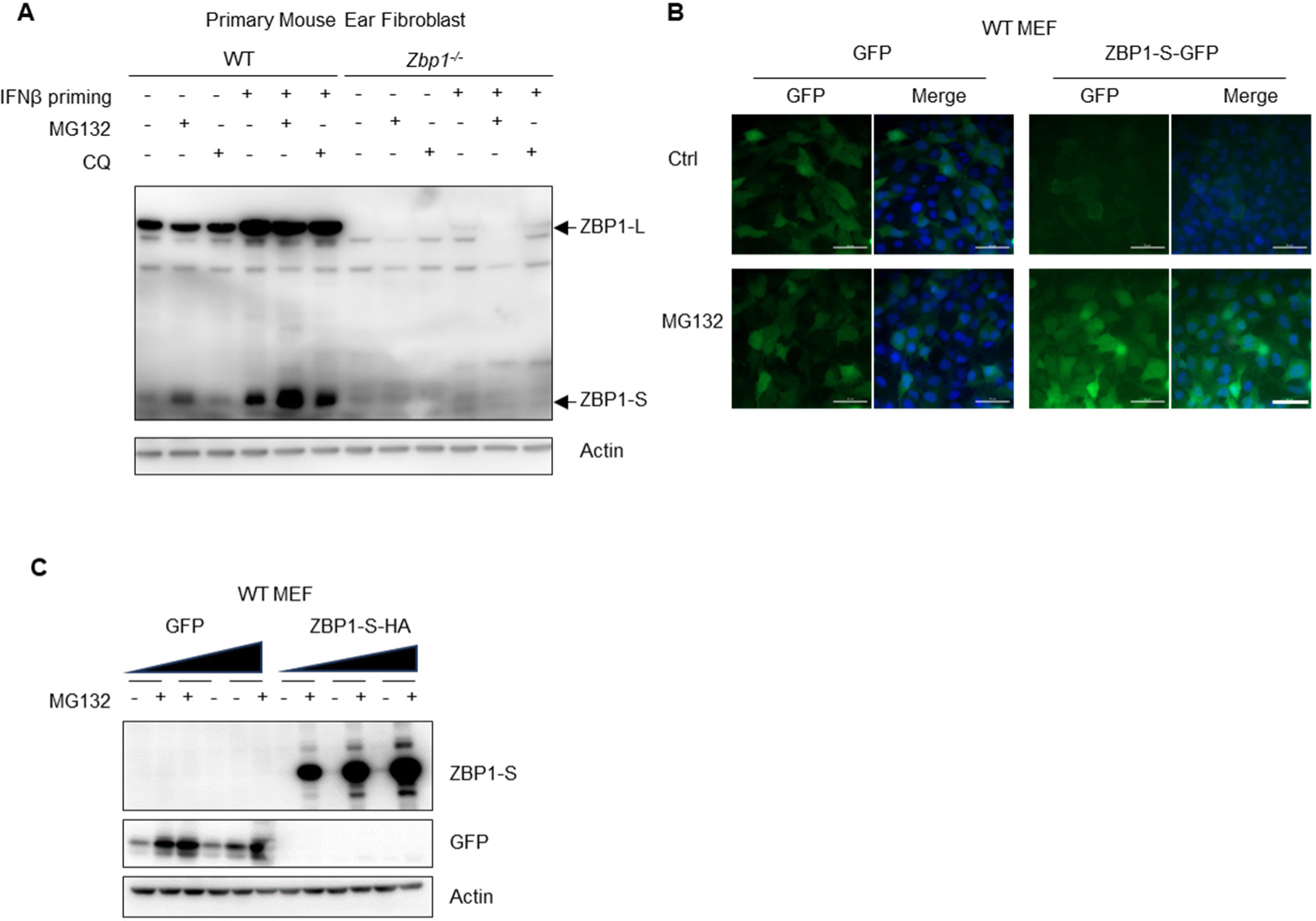
ZBP1-S protein is sensitive to proteasome-mediated degradation. **A.** Protein level of ZBP1-L and ZBP1-S were analyzed by immunoblotting after stimulation of IFNβ (10 ng/ml) for 24 h in wild-type (WT) or *Zbp1^-/-^* primary mouse ear fibroblasts. Proteasome inhibitor MG132 (10 μM) and autophagy inhibitor chloroquine (CQ, 10 μM) was treated. **B.** ZBP1-S-GFP and GFP were overexpressed in WT MEF cells. Fluorescence images were obtained after the cells were treated by MG132 (10 μM) for 3 h. Scale bar, 50 μm. **C.** The same cells in (B) were treated by MG132 (10 μM) for 12 h. The cell lysates were analyzed by immunoblotting to detect the proteins as indicated.

**Figure S2.**
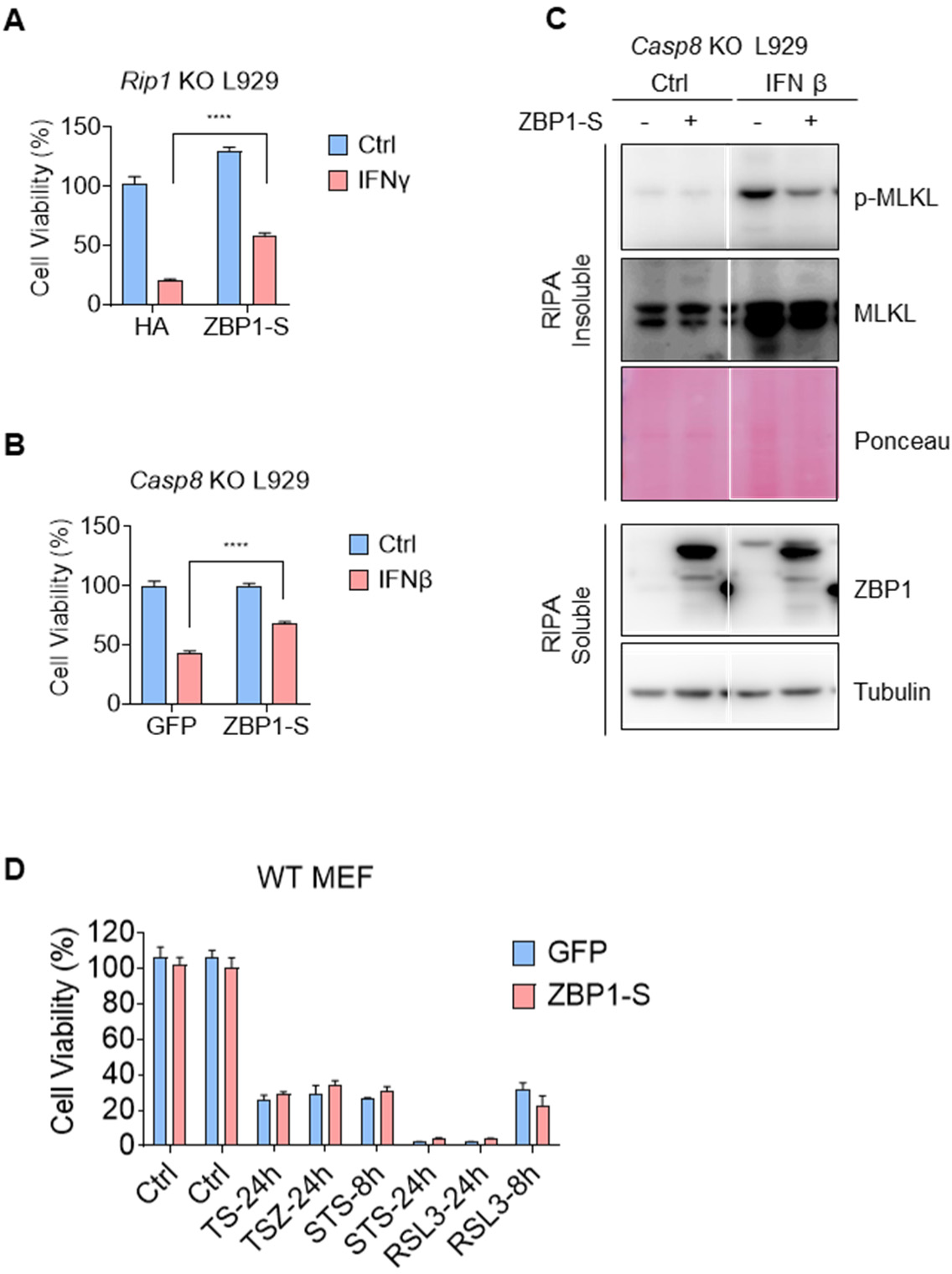
ZBP1-S specifically inhibits ZBP1-dependent cell death. **A.** ZBP1-S-HA and HA-vector were overexpressed in *Rip1* KO L929 cells. Cell survival (%) was measured after treatment of IFNγ (10 ng/ml) for 36 h. Data are shown as mean ± s.e.m. of triplicates. **B.** ZBP1-S-GFP and GFP were overexpressed in *Casp8* KO L929 cells. Cell survival (%) was measured after treatment of IFNβ (10 ng/ml) for 24 h. Data are shown as mean ± s.e.m. of triplicates. **C.** The same cells as in (B) were treated by IFNβ (10 ng/ml) for 24 h. The cell lysates were analyzed by immunoblotting to detect the proteins as indicated. **D.** ZBP1-S-GFP and GFP were overexpressed in WT MEF cells. Cell survival (%) was measured after treatment of TNFα+SMAC mimic (TS), TNFα+SMAC mimic +Z-VAD (TSZ), STS and RSL3. Data are shown as mean ± s.e.m. of triplicates.

**Figure S3.**
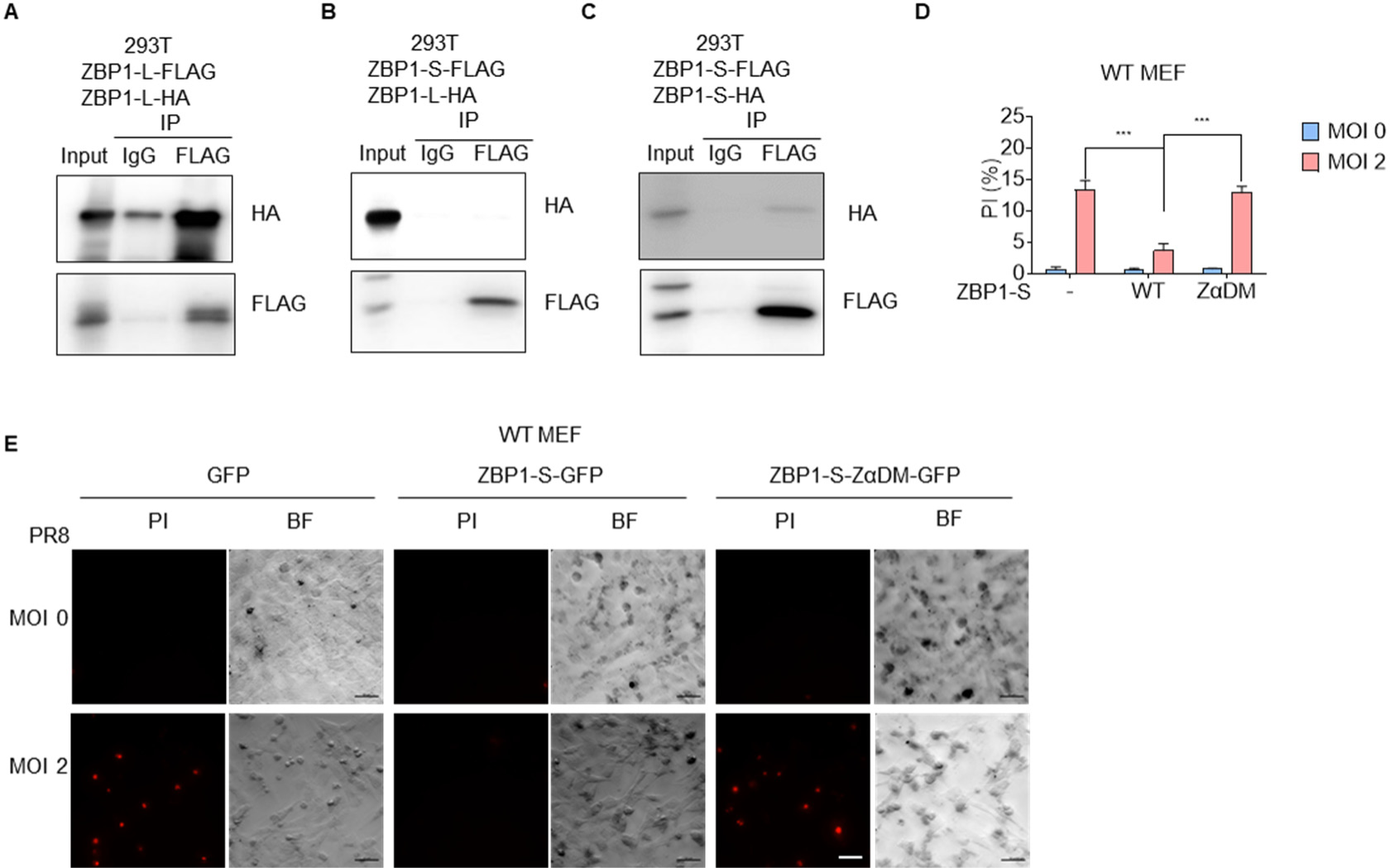
ZBP1-S inhibits ZBP1-dependent cell death by its Z*α* domain. **A.** Immunoprecipitation with anti-FLAG or anti-IgG antibody was performed in 293T cells overexpressed with FLAG-ZBP1-L and HA-ZBP1-L. The cell lysates and immunoprecipitates were analyzed by immunoblotting with antibodies as indicated. **B.** Immunoprecipitation with anti-FLAG or anti-IgG antibody was performed in 293T cells overexpressed with FLAG-ZBP1-S and HA-ZBP1-L. The cell lysates and immunoprecipitates were analyzed by immunoblotting with antibodies as indicated. **C.** Immunoprecipitation with anti-FLAG or anti-IgG antibody was performed in 293T cells overexpressed with FLAG-ZBP1-S and HA-ZBP1-S. The cell lysates and immunoprecipitates were analyzed by immunoblotting with antibodies as indicated. **D.** GFP, ZBP1-S-GFP and ZBP1-S-ZαDM-GFP were overexpressed in WT MEF cells. The strain of Influenza A virus and the MOI were indicated. Percentage of PI positive cells were measured after infection of influenza virus A for 16 h. Data are shown as mean ± s.e.m. of triplicates. ZBP1-S-ZαDM-GFP (ZBP1-S-Zα1& Zα1 domains Double Mutation), the mutation results in disability of Z-nucleic acid binding. Scale bar, 50 μm. **E.** PI images were obtained in the same cell as in (D) after infection of influenza virus A for 16 h.

**Figure S4.**
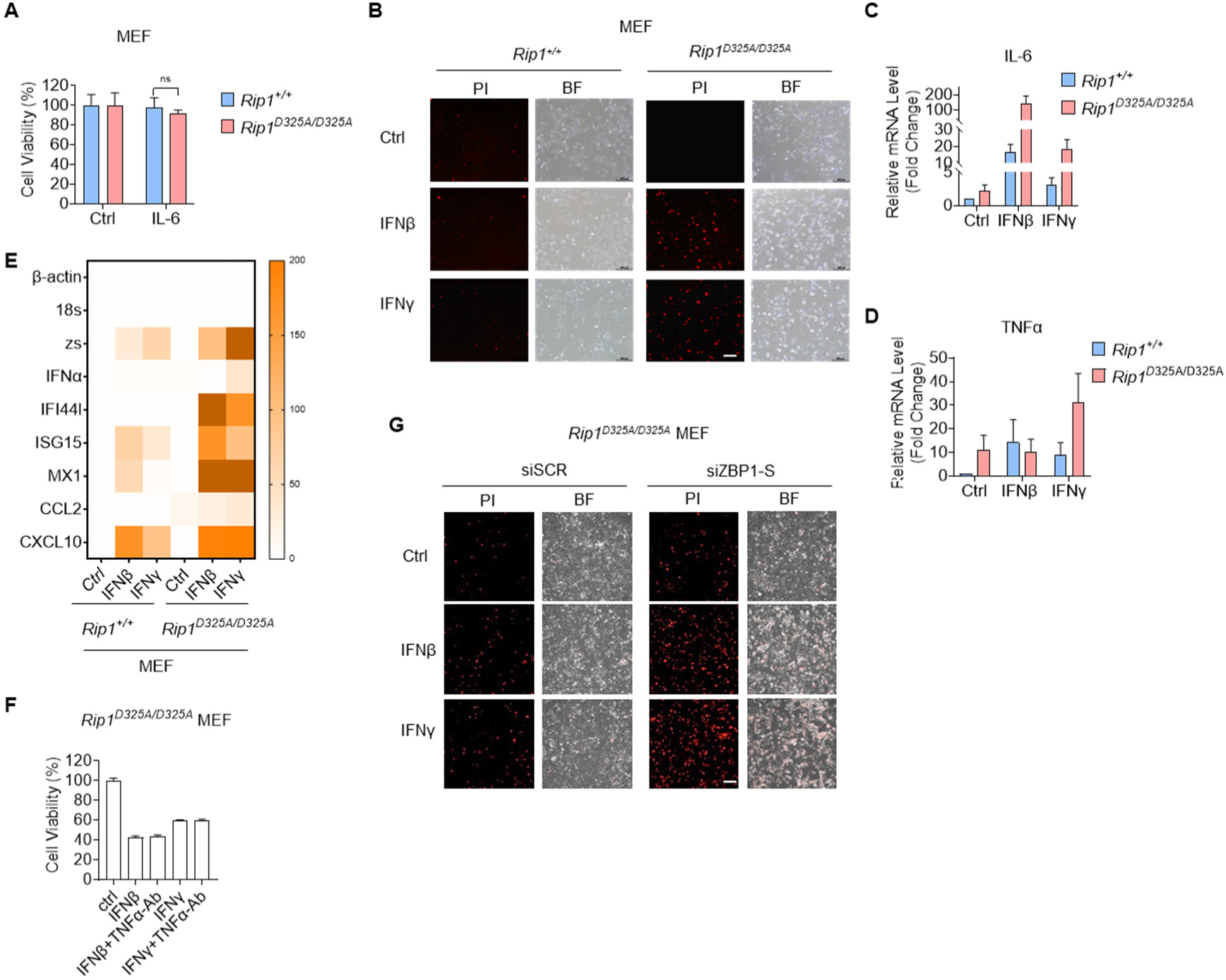
ZBP1-S prevents the autoactivation of ZBP1 in *Rip1^D^*^325^*^A/D^*^325^*^A^* cells. **A.** E1A+Ras-transformed *Rip1^D^*^325^*^A/D^*^325^*^A^* and *Rip1^+/+^* MEFs were treated with IL-6 for 36 h. Cell survival (%) was measured. Data are shown as mean ± s.e.m. of triplicates. **B.** E1A+Ras-transformed *Rip1^D^*^325^*^A/D^*^325^*^A^* and *Rip1^+/+^* MEFs were treated with IFNβ (20 ng/ml) and IFNγ (20 ng/ml) for 36 h. PI images were obtained to detect cell death. Scale bar, 150 μm. **C.** mRNA level of IL-6 was detected after stimulation of IFNβ (20 ng/ml) and IFNγ (20 ng/ml) for 48 h in E1A+Ras-transformed *Rip1^D^*^325^*^A/D^*^325^*^A^* and *Rip1^+/+^* MEFs. Data are shown as mean ± s.e.m. of triplicates. **D.** mRNA level of TNFα was detected after treatment of IFNβ (20 ng/ml) and IFNγ (20 ng/ml) for 48 h in E1A+Ras-transformed *Rip1^D^*^325^*^A/D^*^325^*^A^* and *Rip1^+/+^* MEFs. Data are shown as mean ± s.e.m. of triplicates. **E.** Transcriptional changes of genes downstream of type 1 interferon were detected by RT-qPCR after treatment of IFNβ (20 ng/ml) and IFNγ (20 ng/ml) for 48 h in E1A+Ras-transformed *Rip1^D^*^325^*^A/D^*^325^*^A^* and *Rip1^+/+^* MEFs. **F.** Cell survival (%) of *Rip1^D^*^325^*^A/D^*^325^*^A^* MEFs were measured after treatment by IFNβ (20 ng/ml), IFNβ (20 ng/ml) +anti-TNFα, IFNγ (20 ng/ml) and IFNγ (20 ng/ml) +anti-TNFα. Data are shown as mean ± s.e.m. of triplicates. **G.** ZBP1-S was knocked down by siRNA in E1A+Ras-transformed *Rip1^D^*^325^*^A/D^*^325^*^A^* MEFs. PI images of the cells were obtained after treatment of IFNβ (20 ng/ml) and IFNγ (20 ng/ml) for 36 h. Scale bar, 150 μm.

## STAR★METHODS

Detailed methods are provided in the online version of this paper and include the following:

- KEY RESOURCES TABLE
- RESOURCE AVAILABILITY

- Lead contact
- Materials availability
- Data and code availability
- EXPERIMENTAL MODEL AND SUBJECT DETAILS

- **B.** Cell lines
- METHOD DETAILS

- Cell death assay
- Plasmid construction
- Lentivirus preparation and infection
- IAV infection
- Crypts isolation, organoid culture and treatment
- QUANTIFICATION AND STATISTICAL ANALYSIS

## KEY RESOURCES TABLE

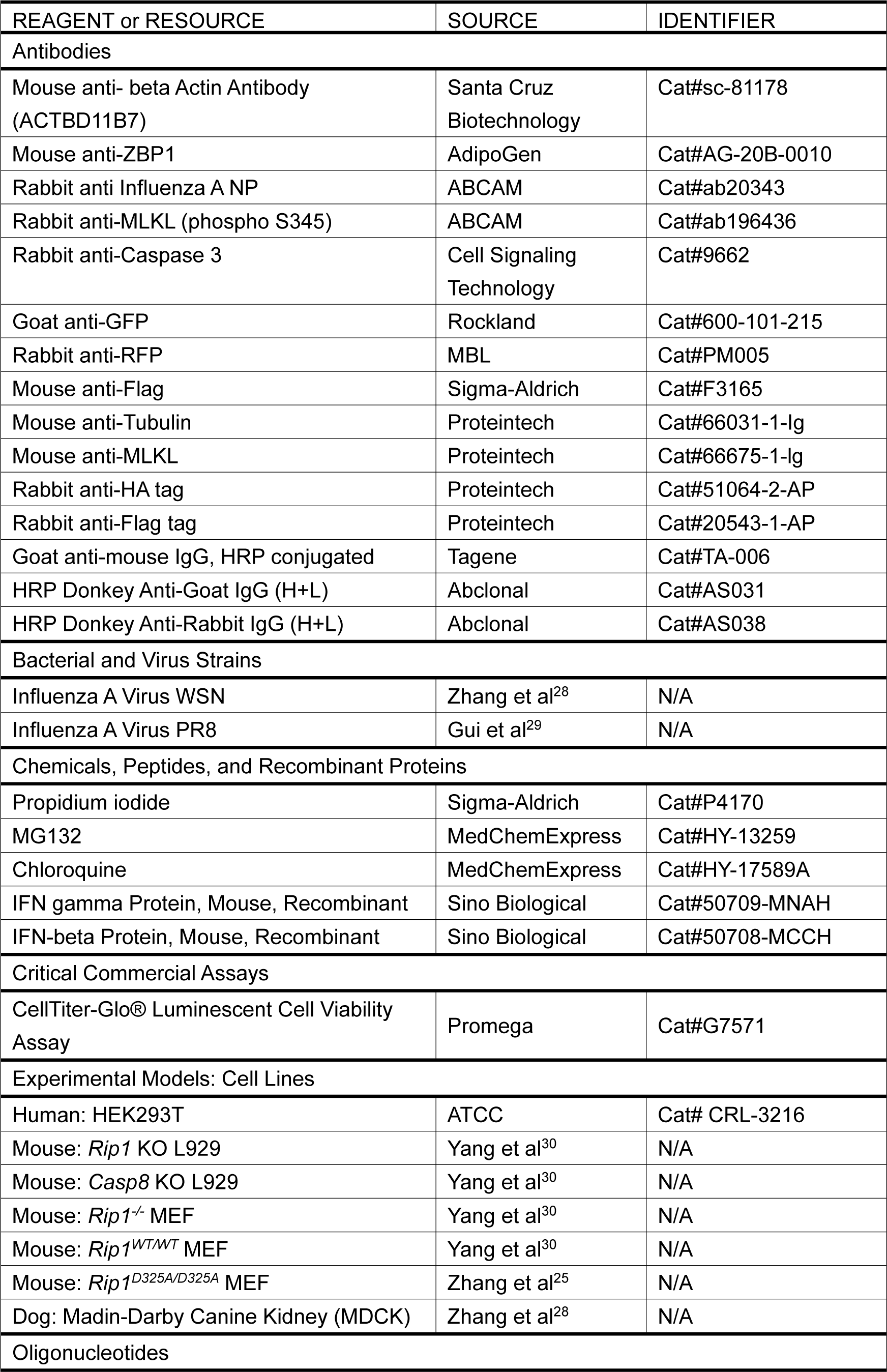
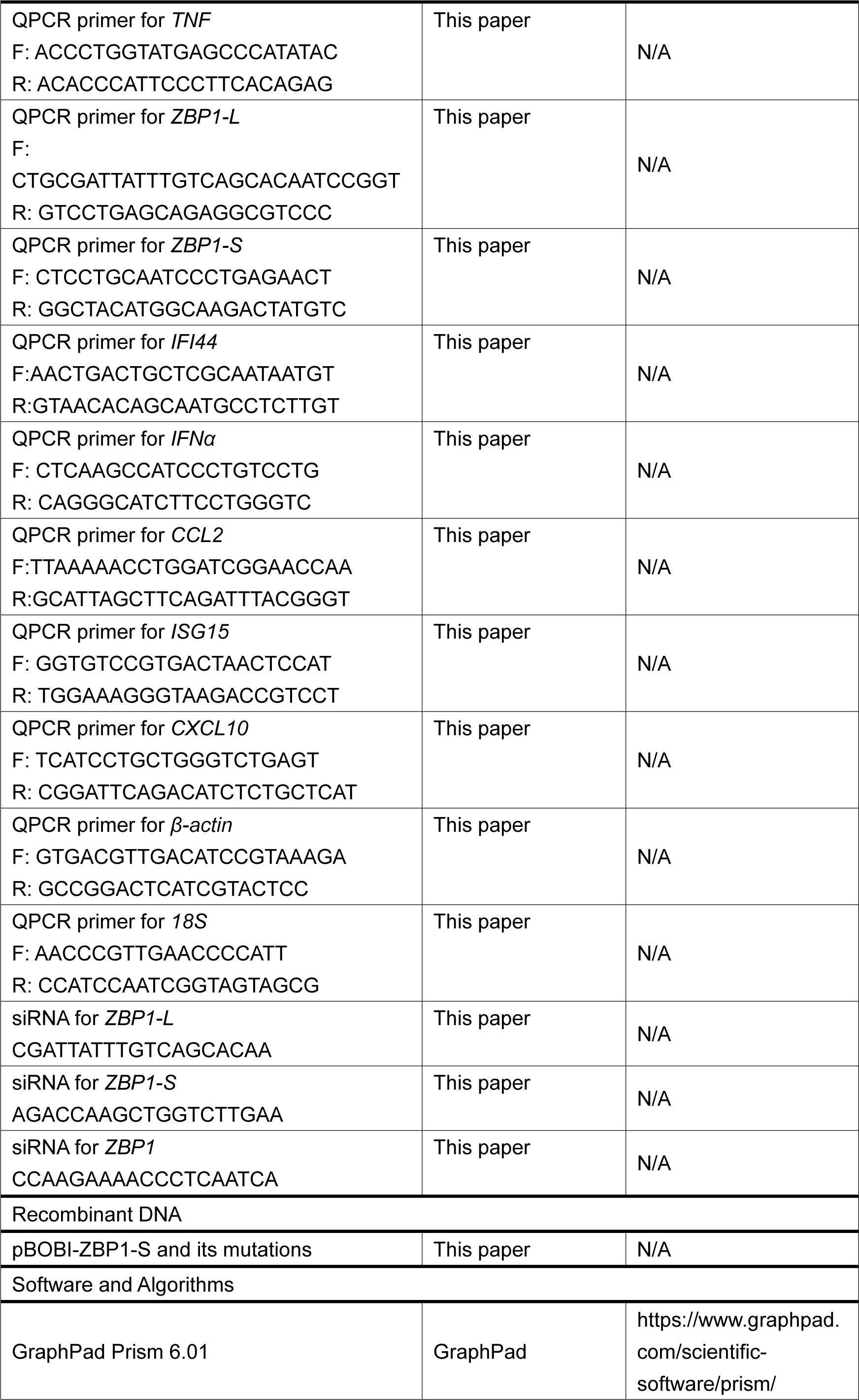
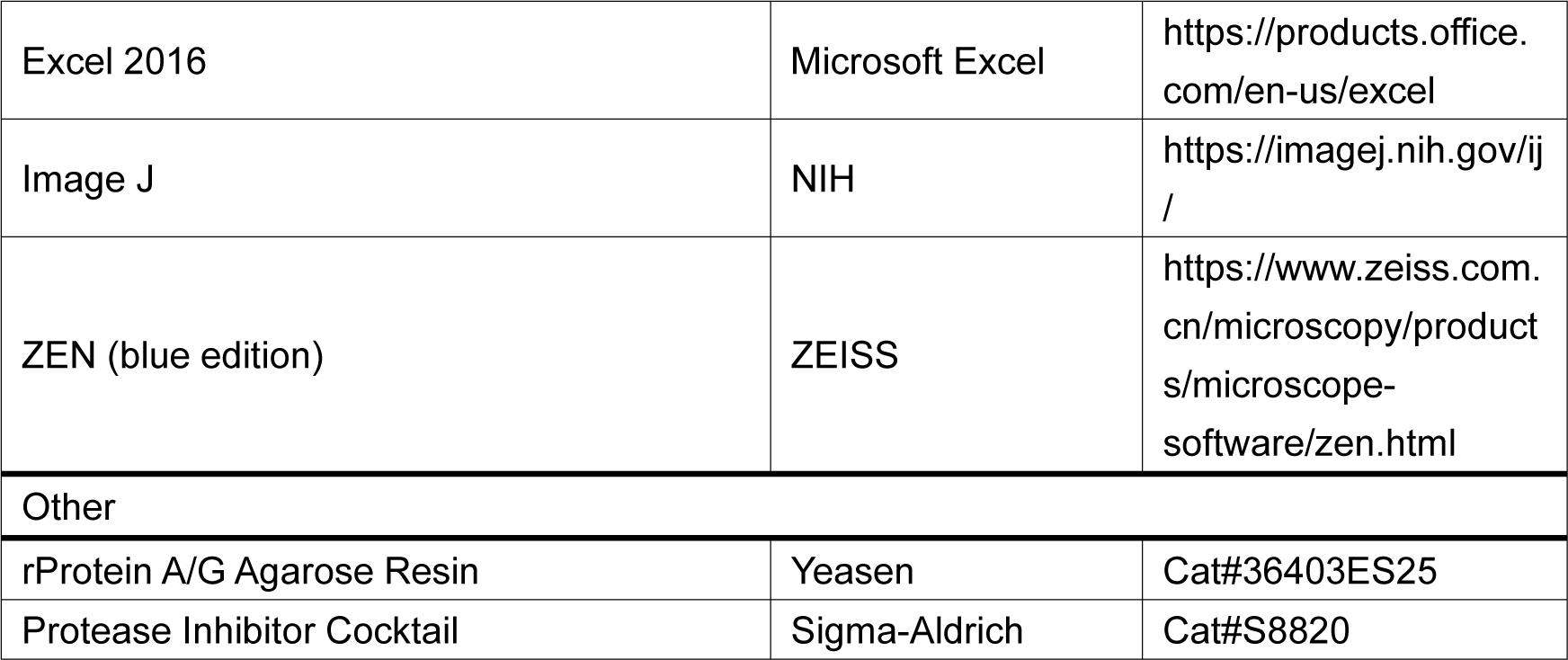

### RESOURCE AVAILABILTY

#### Lead contact

Further information and requests for resources and reagents should be directed to and will be fulfilled by the Lead contact, Zhang-Hua Yang.

#### Materials availability

All plasmids, reagents, cell lines and mouse lines generated in this study are available from the Lead contact.

#### Data and code availability

- Immunoblot images data and microscopy images reported in this paper will be shared by the Lead contact upon request.
- This paper does not report original code.
- Any additional information required to reanalyze the data reported in this paper is available from the Lead contact upon request.

### EXPERIMENTAL MODEL AND SUBJECT DETAILS

#### Cell lines

Mouse fibrosarcoma L929, HEK 293T cells, *Casp8* KO L929*, Rip1* KO L929*, Rip1^-/-^* MEF cells and *Rip1^D^*^325^*^A/D^*^325^*^A^*MEF cells were kindly provided by Prof. Jiahuai Han. Madin-Darby Canine Kidney (MDCK) cells was obtained from Prof. Liang Zhang. Cells were cultured in DMEM (Gibco, USA) supplemented with 10% FBS (vol/vol) (HyClone, USA), 100 IU penicillin (Sangon, China) and 100 mg/ml streptomycin (Sangon, China) at 37 °C in a humidified incubator containing 5% CO2.

## METHOD DETAILS

### Cell death assay

Cell death was analyzed by using a CellTiter-Glo Luminescent Cell Viability Assay Kit (Promega, Madison, WI, USA) according to the manufacturer’s instructions. Briefly, 2 × 104 cells were seeded in 96-well plates with white walls (Nunc). After 12 h, the cells were treated with reagents for the indicated durations. Then, an equal volume of CellTiter-Glo reagent was added to the cell culture medium before the medium was equilibrated to room temperature for 30 min. Cells were shaken for 15 min and rested for 5 min. Luminescence recording was read with a POLARstar Omega (BMG Labtech, Durham, NC, USA).

### Plasmid construction

The full-length sequences of ZBP1 were amplified from cDNA. ZBP1 mutations were introduced by two-round PCR. All these DNA fragments were cloned into pBOBI lentiviral vectors with Flag/HA/GFP tags. The Exo III-assisted ligation-independent cloning method was used for subcloning. All plasmids were verified by DNA sequencing.

### Lentivirus preparation and infection

293T cells were cotransfected with pBOBI constructs and lentiviral packaging plasmids (PMD2/PSPAX) by PEI. Fresh medium was replaced after 12 h. The lentivirus-containing supernatant was collected 36 h later and used for infection. Target cells were infected with lentivirus in the presence of 10 μg/ml polybrene and then centrifuged at 2500rpm for 30 min. The infectious medium was changed 12 h later.

### IAV infection

IAV strains WSN was obtained from Prof. Liang Zhang. Virus titers were determined by plaque assay on Madin-Darby Canine Kidney (MDCK) cells. For cell-culture experiments, near-confluent monolayers of cells were infected with virus in serum-free DMEM for 1 hr in cell incubator, with a gentle rocking every 10 min. Following infection, the inoculum was removed and replaced with growth medium. In conditions involving small-molecule inhibitors. Meanwhile, the Propidium Iodide (PI) was added into medium. Images were analyzed by ZEISS software and.

### Crypts isolation, organoid culture and treatment

The methods of Crypts isolation, organoid culture, and drugs treatment were performed as described previously^5^.

## QUANTIFICATION AND STATISTICAL ANALYSIS

Statistical analysis was performed with Prism software (GraphPad Software). Data are presented as the means ± s.e.m. A two-tailed Student’s t-test was used to compare differences between treated groups and their paired controls.

